# Binucleated human hepatocytes arise through loss of membrane anchorage to the midbody during endomitosis

**DOI:** 10.1101/2023.04.13.536716

**Authors:** Gabriella Darmasaputra, Susana M. Chuva de Sousa Lopes, Hans Clevers, Matilde Galli

## Abstract

Many plant and animal cells transition from canonical to non-canonical cell cycles during development, resulting in the formation of polyploid cells. Two types of non-canonical cell cycles exist: endoreplication, where cells increase their DNA content without entering M phase, and endomitosis, where cells enter M phase but exit prematurely. Although endoreplication has been extensively studied in plants and insects, much less is known on the regulation of endomitosis, which is the most common mode of polyploidization in mammals. In this study, we use fetal-derived human hepatocyte organoids (Hep-Org), to investigate how human hepatocytes initiate and execute endomitosis. We find that cells in endomitosis M phase have normal mitotic timings, but lose membrane anchorage to the midbody during cytokinesis, resulting in regression of the cytokinetic furrow and formation of binucleate cells. Using immunofluorescence, we find that three cortical anchoring proteins, RacGAP1, anillin, and citron kinase (CIT-K), lose their association with the cell cortex during cytokinetic regression. Moreover, reduction of WNT activity by withdrawal of CHIR99021, a GSK3 inhibitor, from the culturing medium increases the percentage of binucleated cells in Hep-Orgs. This effect is lost in organoids with mutations in the atypical E2F proteins, E2F7 and E2F8, which have been implicated in binucleation of rodent hepatocytes. Together, our results identify how human hepatocytes inhibit cell division in endomitosis, and highlight an evolutionary recurrent mechanism to initiate non-canonical cell cycles in mammals.

## Introduction

Polyploid cells, which contain more than two pairs of homologous chromosomes, are found in many tissues of diploid species, including mammals. Somatic polyploidization occurs during defined moments in development, and is thought to be crucial for increases in metabolic output^1,2^, cell and organism size^3^, and to maintain specialized cell functions^4–6^. Polyploidization has also been shown to have a protective role against genotoxic stress, by buffering the effect of detrimental genetic aberrations^7–9^. In humans, polyploid cells arise in tissues such as the liver, pancreas, mammary glands, placenta, and bone marrow^10–12^. In these tissues, cells become polyploid by undergoing non-canonical cell cycles, so-called endocycles, in which they replicate their DNA but do not divide. Two types of endocycles have been described that give rise to polyploidy: endoreplication and endomitosis. In endoreplication, cells alternate between S and G phases, duplicating their genomic DNA without ever entering M phase. In endomitosis, cells enter M phase but exit prematurely, before completing cell division. Depending on the timing of M-phase exit, daughter cells can either be mononucleated or binucleated.

An outstanding question in cell-cycle research is how cells transition from canonical to non-canonical cell cycles. Most prior work has focused on the transition from canonical cycles to endoreplication, which has been extensively studied in plants and insects and relies on downregulation of M-CDK activity^11^. By suppressing M-CDK, cells cannot enter M phase, and this is sufficient to trigger endoreplicative cycles in animals and yeast^5,13–15^. On the other hand, less is known on how cells transition to endomitosis cycles. In endomitosis, M-CDK needs to become active to initiate M phase, yet cytokinesis needs to be inhibited to block cell division. The mechanisms by which endomitotic cells inhibit cytokinesis differ per cell type and even during different stages of development of the same cell type. For example, during M phase of the first endomitosis cycle of megakaryocytes, cells undergo cleavage furrowing followed by regression of the furrow, whereas there is no cytokinetic ingression in subsequent endomitosis cycles^16,17^. Also in other endomitotic cells, such as mouse cardiomyocytes^18,19^, hepatocytes^20–23^, and mammary cells^1^ cell division is inhibited at different stages of M phase. Despite different mechanisms to inhibit cell division between different endomitotic cell types, downregulation or inhibition of key cytokinesis regulators seems to be a common feature.

Although most polyploid cells are considered terminally differentiated and post-mitotic, a few types of polyploid cells have been found to remain proliferative. In *Drosophila*, rectal papillae cells become polyploid through endoreplication, but can switch back to mitotic divisions during later development or upon injury^24,25^. Similarly, in regenerating mouse livers, polyploid hepatocytes are able to divide and revert their ploidy^26–28^. The ability of cells to switch back from non-canonical cycles to canonical cycles suggest that the inhibition of cell division is reversible. This phenomenon is likely important for the regenerative capacity of the liver, as it has been shown that polyploid hepatocytes contribute to liver regeneration in mice^27–29^. Thus, deciphering the molecular mechanisms by which hepatocytes switch between canonical and non-canonical cycles will allow a deeper understanding of how tissues modulate the percentages of polyploid cells, and may identify novel factors that control regeneration.

The mammalian liver is an attractive model to study the regulation of non-canonical cell cycles, as mature hepatocytes have been reported to perform both canonical and endomitotic cell cycles throughout their lifetime. In humans, hepatocytes range in ploidy from 2N to 8N, and up to 20% are binucleated^30–33^. In a healthy liver, less than 0.01% of hepatocytes are actively cycling^34,35^. However, upon injury, hepatocytes are able to proliferate and regrow the organ to an equivalent of the original size. Proliferating mouse hepatocytes require Wnt/β-catenin activity^36^, but high Wnt/β-catenin activity by itself does not induce proliferation^36,37^. This is in contrast with many other mammalian epithelia where the degree of proliferation strongly correlates with Wnt activity^38^, suggesting that the Wnt/β-catenin axis may have an unconventional role in controlling hepatocyte cell cycles.

Hepatocyte endomitosis has thus far mostly been studied in rodents, and it is currently unknown how human hepatocytes undergo endomitosis. In mouse livers, hepatocytes undergo canonical cell cycles during embryogenesis, but transition to endomitosis cycles during postnatal development^39^. Here, the activity of the atypical E2F transcription factors E2F7 and E2F8 drives endomitosis cycles by repressing cytokinesis genes^23^. It is unclear whether WNT/ß-catenin signaling and E2F7/8 regulate endomitosis in human hepatocytes. Investigation of hepatocyte canonical and non-canonical cycles *in vitro* is not trivial, as primary hepatocyte cultures are limited in their proliferative capacity^40^ and require a three-dimensional structure for proper polarization and proliferation^41,42^. Immortalized non-cancerous hepatocyte cell lines are also unsuitable for cell-cycle analyses as they often have impaired physiology^43^. In this study, we use fetal tissue-derived human hepatocyte organoids (Hep-Org)^44^ as a model to study the regulation of endomitosis. Hep-Orgs can be cultured long-term and be genetically modified, allowing integration of fluorescently-tagged markers and live imaging of canonical and endomitosis M phases^45^. We find that human hepatocytes undergoing endomitosis initiate cytokinesis with normal timing and morphology, but regress their cytokinetic furrows during late M phase. Our immunofluorescence analyses of cells undergoing regression suggest that endomitotic cells have defects in tethering the midbody to the cell cortex. We also find that inhibition of WNT signaling increases the percentage of binucleated cells, a process that is dependent on E2F7 and E2F8. Together, our work reveals a novel mechanism for the inhibition of cytokinesis during endomitosis M phase, and provides support for an evolutionary conserved function of WNT and E2F7/E2F8 in the transition to non-canonical cell cycles.

## Results

### Hep-Orgs represent a model to study canonical and endomitosis M phases in vitro

To determine whether Hep-Orgs can be used as a model to study the regulation of endomitosis, we first determined the percentage of binucleated cells in three different liver organoid lines. We generated organoid strains expressing a nuclear GFP-NLS marker and used CellMask Orange dye to mark the cell membrane (**Fig. 1A**). Consistent with published observations on the frequencies of binucleated hepatocytes in fetal human hepatocytes^31^, we observed an average of 5% and 15% binucleated cells in two independent Hep-Org lines (**Fig. 1B**). In contrast, binucleated cells are rarely found (0.8%) in cholangiocyte organoids (Chol-Org) (**Fig. 1B**). The percentage of binucleated cells in each organoid ranged between 0-32% and did not show significant differences between organoids of different sizes (**Fig. 1C**).

**Figure 1.**
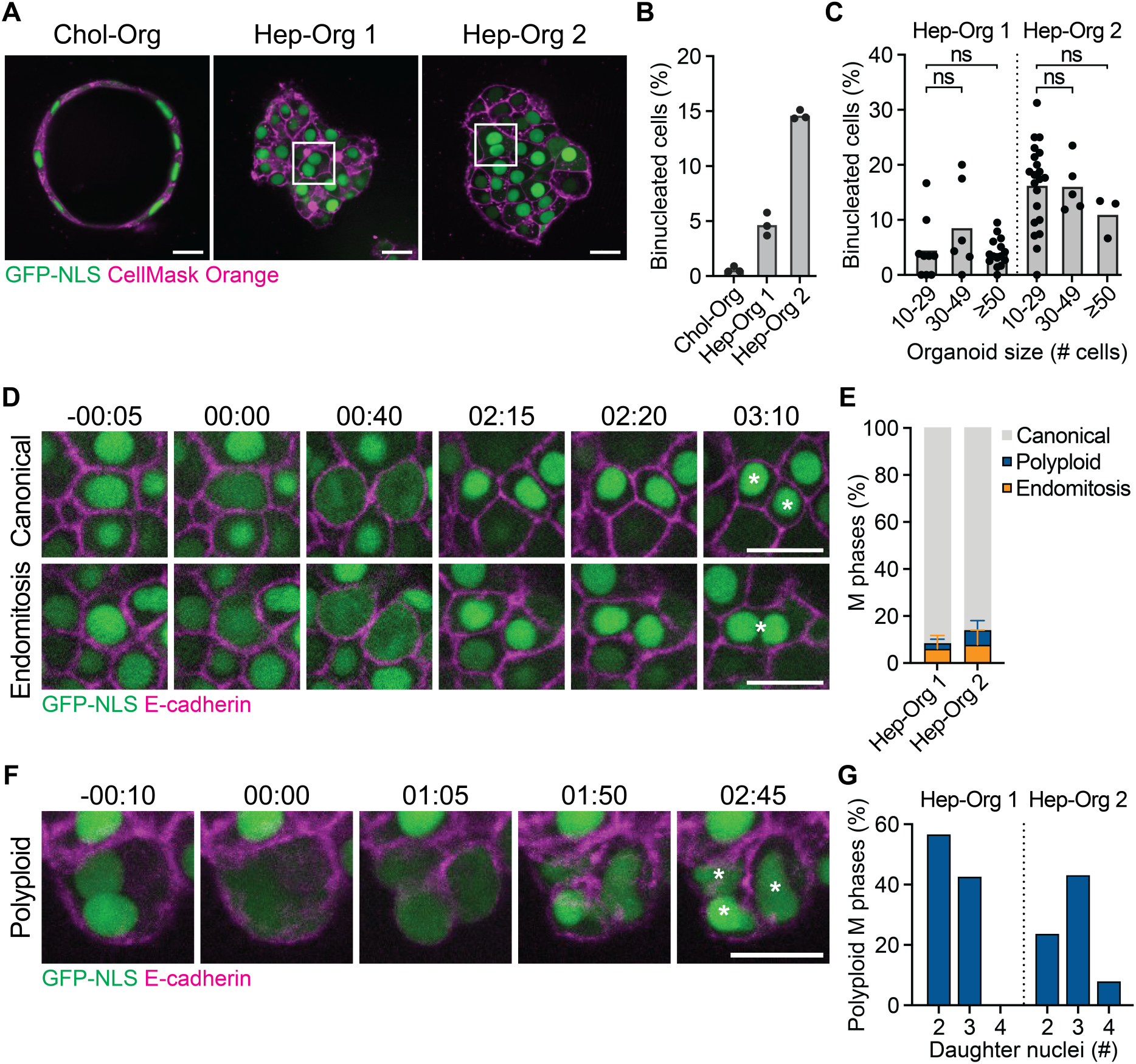
Human hepatocytes undergo canonical, endomitosis and polyploid M phases in Hep-Orgs. **A**) Representative images of Chol-Org and Hep-Org lines expressing GFP-NLS (shown in green) and stained with CellMask Orange (shown in magenta) to mark nuclei and membranes, respectively. Images show one plane in the center of each organoid. Examples of binucleated cells are indicated in the white squares. **B**) Percentage of binucleated cells in Chol-Org and Hep-Org lines. Columns depict mean percentages from 3 biologically independent experiments (200-300 cells analyzed per experiment). **C**) Percentage of binucleated cells per organoid plotted against organoid size (n = 3 experiments, 20-30 organoids analyzed per Hep-Org line). Each dot represents one organoid, with the mean depicted in column (ns = not significant, Student’s *t*-test). **D**) Stills from live-imaging of GFP-NLS/E-cadherin-tdTom Hep-Org 1 line showing canonical (top) and endomitosis (bottom) M phase. Time is relative to nuclear envelope breakdown (NEB) in hh:mm. White asterisks mark daughter cells. **E**) Percentage of types of M phases observed during live-imaging experiments of GFP-NLS/E-cad-tdTom Hep-Org 1 line and Tubulin-mNeon Hep-Org 2 line (n = 5 experiments, >175 events analyzed per line). Error bars represent standard deviation. **F**) Stills from live-imaging of GFP-NLS/E-cadherin-tdTom Hep-Org 1 line showing polyploid M phase. Time is relative to NEB in hh:mm. White asterisks mark daughter cells. **G**) Percentage of polyploid M phases segregating their DNA content to 2, 3, or 4 daughter nuclei (n = 5 experiments, >7 events per line). Scale bars in A, D, and, F represent 50 µm.

To visualize endomitosis in Hep-Org, we performed long-term live imaging of Hep-Orgs expressing a nuclear GFP marker and endogenously tagged E-cadherin-tdTomato to mark cell membranes (referred to as GFP-NLS/E-cadherin-tdTom), or endogenously tagged β-tubulin-mNeonGreen to visualize the mitotic spindle (referred to as Tubulin-mNeon) (**Fig 1D, F**). Cells undergoing M phase were identified based on either the loss of a confined GFP-NLS signal during nuclear envelope breakdown (NEB) in the GFP-NLS/E-cadherin-tdTom line, or by the appearance of spindle poles in the Tubulin-mNeon line (**Fig 1D, 1F, 2A**). We categorized different types of M phases, depending on the number of nuclei in the mother and daughter cells. We found that 90% of M phases in both Hep-Org lines were canonical, where a mononucleated mother cell gives rise to two mononucleated daughter cells (**Fig. 1D**, top panel). In contrast, 5-8% of M phases showed a mononucleated mother cell giving rise to one binucleated daughter cell, which we classify as endomitosis M phases (**Fig. 1D**, bottom panel).

Finally, we also observed binucleated cells entering M phase (**Fig. 1F**), showing that polyploid cells in Hep-Org continue cycling, as has been shown previously^45^. These divisions, which we refer to as polyploid M phases, give rise to either two, three, or four daughter nuclei (**Fig 1G**). Taken together, we observe canonical, endomitosis, and polyploid M phases in Hep-Orgs, rendering them an attractive model to study the transition between canonical and non-canonical cell cycles in human hepatocytes *in vitro*.

### Cells undergoing endomitosis M phase have normal mitotic timings but regress their cytokinetic furrow during late cytokinesis

Previous studies on endomitosis in mouse megakaryocytes, hepatocytes, and cardiomyocytes have identified distinct mechanisms to inhibit cytokinesis in endomitosis M phase^16–19,21,22,46^. To determine how human hepatocytes inhibit cell division during endomitosis, we performed live-imaging on Hep-Org lines with endogenously tagged β-tubulin-mNeonGreen and E-cadherin-tdTomato (from here on Tubulin-mNeon/E-cadherin-tdTom) to visualize the formation of the mitotic spindle and characterize cytokinetic events. Similar to canonical M phases, we observed central spindle assembly, furrow ingression, and midbody formation in all cases of endomitosis (**Fig. 2A**, n = 18). However, during late stages of endomitosis M phase, the cell membrane disconnects from the midbody and regresses. Despite being disconnected from the cell membrane, midbody structures remain stable after cytokinetic regression, and undergo normal severing during M-phase exit (**Fig. 2A**).

**Figure 2.**
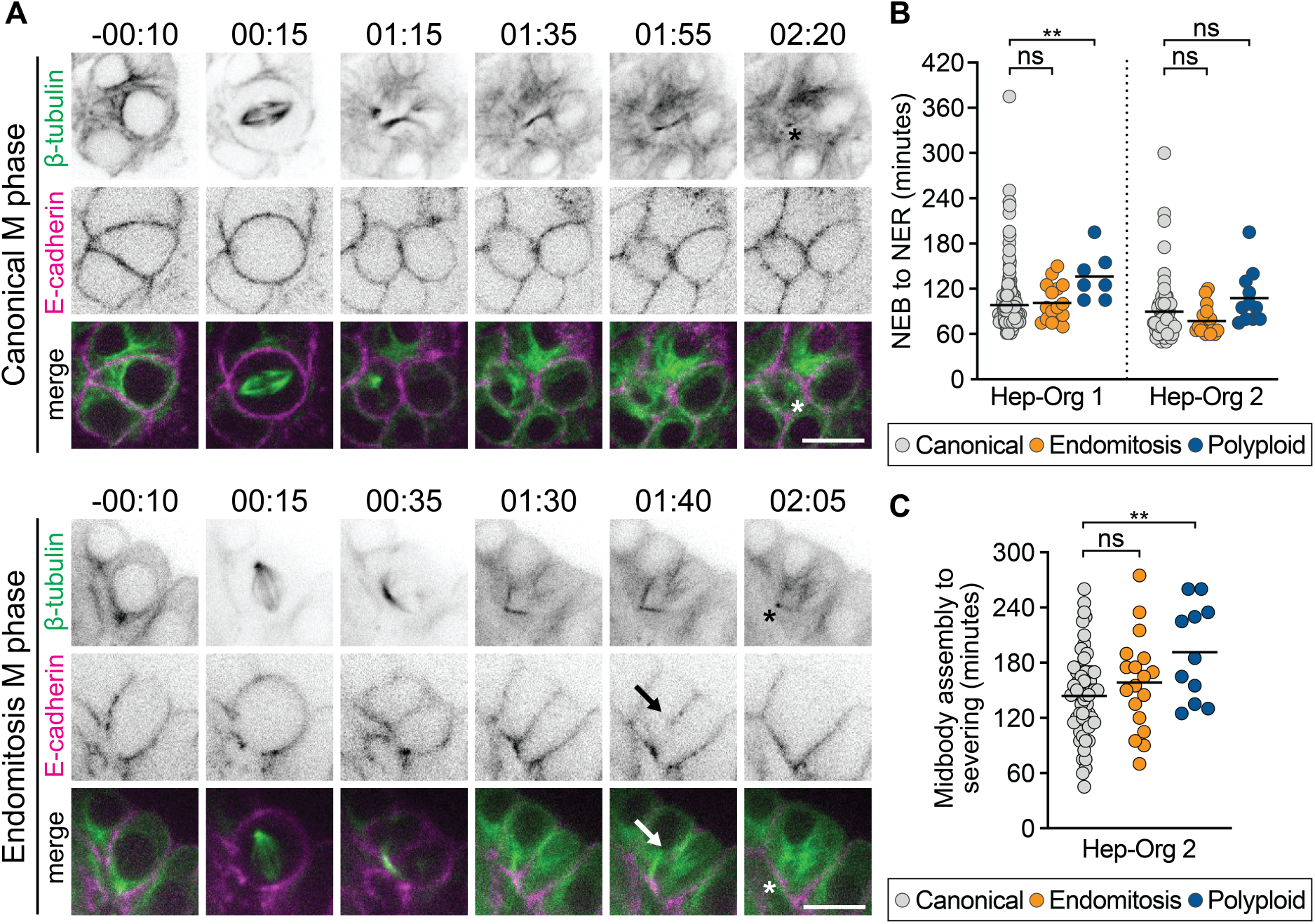
Cells undergoing endomitosis have normal mitotic timings but regress their cytokinetic furrow during late M phase. **A**) Representative stills from live-imaging of Tubulin-mNeon/E-cadherin-tdTom Hep-Org 2 line showing canonical (top) and endomitosis (bottom) M phases. Stills show formation of central spindle in both canonical mitosis and endomitosis, with subsequent membrane regression in endomitosis (marked with arrow) and midbody severing (marked with asterisk). Time is relative to NEB in hh:mm. Scale bars represent 50 µm. Panels 3-6 showing β-tubulin are maximum projections of 2 z-slices. **B**) Duration of NEB to NER in minutes for canonical, endomitosis and polyploid M phases in Hep-Org 1 line with GFP-NLS/E-cadherin-tdTom and Hep-Org 2 line with Tubulin-mNeon. Individual measurements are shown for canonical (Hep-Org 1, n = 204 events; Hep-Org 2, n = 62 events), endomitosis (Hep-Org 1, n = 16 events; Hep-Org 2, n = 18 events), and polyploid (Hep-Org 1, n = 7 events; Hep-Org 2, n = 11 events) M phases. Black bars indicate mean (ns = not significant, \*\**p*<0.01, Student’s *t*-test). **C**) Duration from midbody assembly to disassembly in minutes for canonical (n = 62 events), endomitosis (n = 18 events) and polyploid (n = 11 events) M phases in Hep-Org 2 line with Tubulin-mNeon. Individual measurements are shown with mean (black bar) (\*\**p*<0.01, Student’s *t*-test).

To determine whether there are any differences in mitotic progression between cells undergoing endomitosis or canonical M phases, we compared mitotic timings using the Tubulin-mNeon Hep-Org 2 line. We found that the duration from NEB to nuclear envelope reformation (NER) is very similar between canonical and endomitosis M phases (**Fig. 2B**). Furthermore, despite membrane regression, there is no difference in timing between midbody formation and disassembly in endomitosis compared to canonical M phases (**Fig. 2C**). For polyploid M phases, NEB–NER duration is 1.5-fold higher than canonical and endomitosis M phases. This is not unexpected, considering these cells contain twice as many chromosomes and centrosomes, which will likely delay their mitotic progression. Taken together, this data shows that canonical and endomitosis M phases in human hepatocytes are very similar during early stages of mitosis, but that endomitotic cells inhibit their division at a late step of cytokinesis.

### Cells undergoing endomitosis lose membrane anchoring to the midbody

To understand the underlying cause of membrane regression during endomitosis, we further examined cytokinetic events, zooming in on cytokinetic furrows and midbody structures using immunofluorescent (IF) staining in the E-cadherin-tdTom Hep-Org 1 line. We first determined whether we could identify endomitosis M phases in fixed Hep-Orgs. When looking at static images, ingression of the cleavage furrow during early telophase looks similar to a regressed membrane in endomitosis, making it difficult to distinguish the two by solely looking at the membrane. Nonetheless, by staining for α-tubulin and co-staining with DAPI, we could identify cells with regressed membrane in late telophase by the presence of a midbody structure and decondensed DNA. In contrast, cells in early telophase have condensed DNA and a broader central spindle structure. When looking at all cells in late telophase, we found that 13 out of 169 cells (7.69%) had regressed membranes, which is in line with the percentage of endomitosis M phases that we found using live-imaging (**Fig. 1E**). Thus, we can use IF staining on fixed Hep-Orgs to further analyze regressed structures during endomitosis M phase.

We first investigated whether endomitotic regression could be a consequence of the activation of an abscission checkpoint. Specifically, presence of chromosome bridges in the cleavage plane activates the abscission checkpoint and can result in cytokinetic regression if the chromosome bridges are not resolved^47^. Therefore, we determined whether we could detect any DNA in the cleavage plane of cells undergoing late telophase in E-cadherin-tdTom Hep-Orgs stained with DAPI and anti-α-tubulin. In 179/179 cases, including 13 cells undergoing endomitotic regression, we could not detect any DNA in the cleavage plane. This makes it unlikely that membrane regression in endomitosis is due to the presence of chromosome bridges.

We next focused on cells that were in the process of membrane regression to understand how cytokinesis is inhibited in endomitosis. During canonical M phase, actomyosin ring contraction generates a cleavage furrow that partitions the mother cell into two. Once ingression is complete, several protein complexes anchor the cell membrane to the microtubules of the midbody to stabilize the cytokinetic furrow^48^. Subsequently, actin filaments of the actomyosin ring disassemble and the cell undergoes abscission, in which the endosomal sorting complex required for transport III (ESCRT-III) splits the plasma membrane, giving rise to two daughter cells. In addition to its essential role in membrane abscission, the ESCRT-III machinery is also required to initiate microtubule severing by targeting the microtubule-severing enzyme Spastin to the midbody^48,49^. Since we did not detect any abnormalities in midbody assembly, furrow ingression, or microtubule severing (see **Fig. 2A**), we wondered whether membrane anchoring is impaired during endomitosis M phases. To investigate this, we performed IF staining of three key membrane anchoring components: RacGAP1, Anillin, and citron Rho-interacting kinase (CIT-K).

We focused our analysis on cells in early telophase and cells in late telophase with either ingressed cleavage furrows or with regressed membranes, the latter reflecting cells in endomitosis. We categorized the localizations of RacGAP1, Anillin, and CIT-K during early and late telophase to determine if there were any abnormalities in regressed structures (**Fig. 3A**). The centralspindlin component RacGAP1 was present at the central spindle in early telophase and localized to the midbody ring in late telophase. RacGAP1 was not present at the membrane during early telophase, but its localization overlapped with the E-cadherin signal in late telophase cells with ingressed cleavage furrows (**Fig. 3B**). In regressed structures, RacGAP1 retained its normal localization to the midbody ring but was no longer associated to the cell membrane (**Fig. 3B**). Similarly, Anillin, which initially localized to the cleavage furrow in early telophase, maintained midbody localization in regressed structures but could no longer be detected at the cell membrane (**Fig. 3C**). Analysis of CIT-K localization revealed two distinct localization patterns on the midbody: either restricted to the midbody ring, as has been described before (**Fig. 3D**)^50–52^, or along the whole midbody (**Fig. S1**). The broad midbody localization was observed in 37% of ingressed late telophase structures and 100% of regressed structures (**Table 1**). Furthermore, CIT-K co-localized with E-cadherin in all early telophase and ingressed structures, but was not found on the cortex in cells undergoing endomitotic regression (**Fig. 3D**, **Table 1**). Taken together, we find that the membrane anchoring proteins RacGAP1, Anillin, and CIT-K are present on the midbodies of cells undergoing endomitosis M phase, but lose their association with the membrane. Furthermore, we find that CIT-K localizes to a broader region on the midbody than has previously been described in cells undergoing cytokinesis, which could indicate an altered or impaired function of CIT-K in endomitosis.

**Figure 3.**
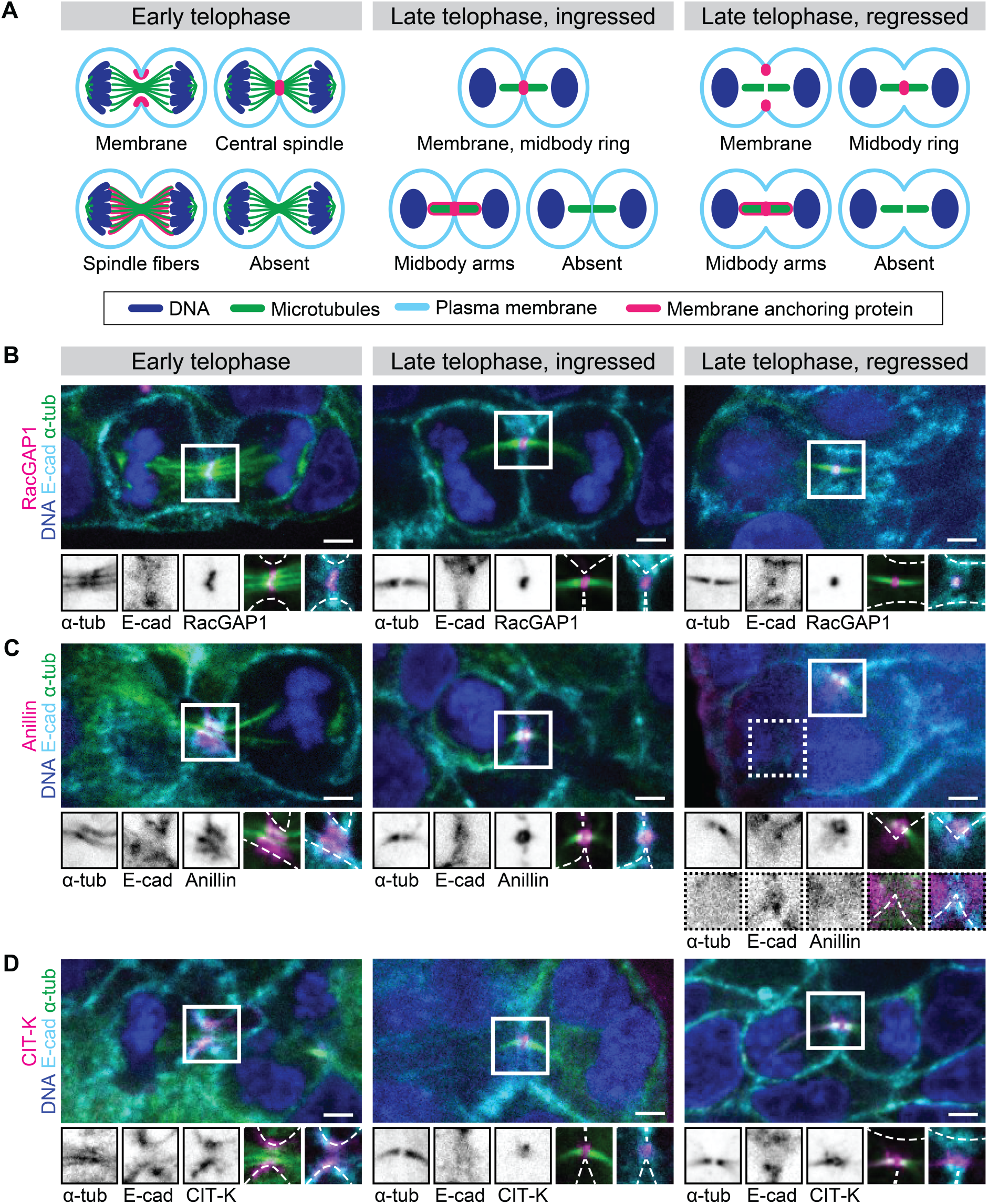
Membrane association of membrane anchoring proteins is lost during cleavage regression in endomitosis. **A**) Schematics illustrating cells in early or late telophase with either ingressed or regressed cleavage furrows, with possible localizations of membrane anchoring proteins. **B–D)** Representative stills of IF experiments showing DAPI, α-tubulin, E-cadherin and membrane anchoring protein RacGAP1 (**B**), Anillin (**C**), or CIT-K (**D**) in early and late telophase. Scale bars represent 3 µm. Close-ups show single or double channel images of marked regions of interest (white box). Dashed lines represent membrane outline. All images show one plane in the middle of the midbody. Quantifications of RacGAP1, Anillin, and CIT-K localizations during the different stages are listed in **Table 1**.

**Table 1.** Quantification of RacGAP1, Anillin, and CIT-K localizations in late telophase with either ingressed or regressed cytokinetic furrows.

| Protein | Localization | Early telophase | Late telophase |  |
| --- | --- | --- | --- | --- |
|  |  |  | Ingressed | Regressed |
| RacGAP1 | Membrane | 0% (0/16) | 100% (67/67) | 0% (0/4) |
|  | Central spindle/<br>Midbody ring | 100% (16/16) | 99% (66/67) | 100% (4/4) |
|  | Spindle fibers/<br>Midbody arms | 0% (0/16) | 0% (0/67) | 0% (0/4) |
|  | Absent | 0% (0/16) | 0% (0/67) | 0% (0/4) |
| Anillin | Membrane | 100% (9/9) | 100% (45/45) | 0% (0/6) |
|  | Central spindle/<br>Midbody ring | 0% (0/9) | 100% (45/45) | 100% (6/6) |
|  | Spindle fibers/<br>Midbody arms | 0% (0/9) | 0% (0/45) | 0% (0/6) |
|  | Absent | 0% (0/9) | 0% (0/45) | 0% (0/6) |
| CIT-K | Membrane | 100% (10/10) | 100% (54/54) | 0% (0/3) |
|  | Central spindle/<br>Midbody ring | 0% (0/10) | 100% (54/54) | 100% (3/3) |
|  | Spindle fibers/<br>Midbody arms | 40% (4/10) | 37% (20/54) | 100% (3/3) |
|  | Absent | 0% (0/10) | 0% (0/54) | 0% (3/3) |

### WNT signaling inhibits binucleation of human hepatocytes in an E2F7/8-dependent manner

Next, we wondered what determines the decision to undergo canonical or endomitosis M phase in hepatocytes. In the liver, hepatocytes are organized in hexagonal lobules. Along the rows of hepatocytes, from the central vein in the middle to the periportal veins at the corners, hepatocytes experience gradients in nutrients, oxygen, and cytokines, and exhibit distinct patterns of gene expression^53–58^. One of these gradients is the WNT gradient, which is secreted from the central vein and has been recently shown to promote canonical mitosis in mouse hepatocytes through Tbx3-mediated repression of E2F7 and E2F8^59^.

Under normal culturing conditions, Hep-Orgs are kept in an active WNT state by the presence of CHIR99021, a GSK-3 inhibitor and potent activator of WNT signaling. To test whether WNT signaling regulates binucleation in human hepatocytes, we removed CHIR99021 from the culture medium for 3 days and quantified the percentage of binucleated cells in Hep-Orgs. Strikingly, we found a two-fold increase in the percentage of binucleated cells after three days of CHIR99021 removal from the medium (**Fig. 4A**). Additionally, Hep-Orgs cultured without CHIR99021 proliferated slower and exhibited some morphological changes, growing in a round shape with a large lumen (**Fig. 4B**). Thus, similar to murine hepatocytes, WNT signaling supports proliferation and inhibits binucleation of human hepatocytes.

**Figure 4.**
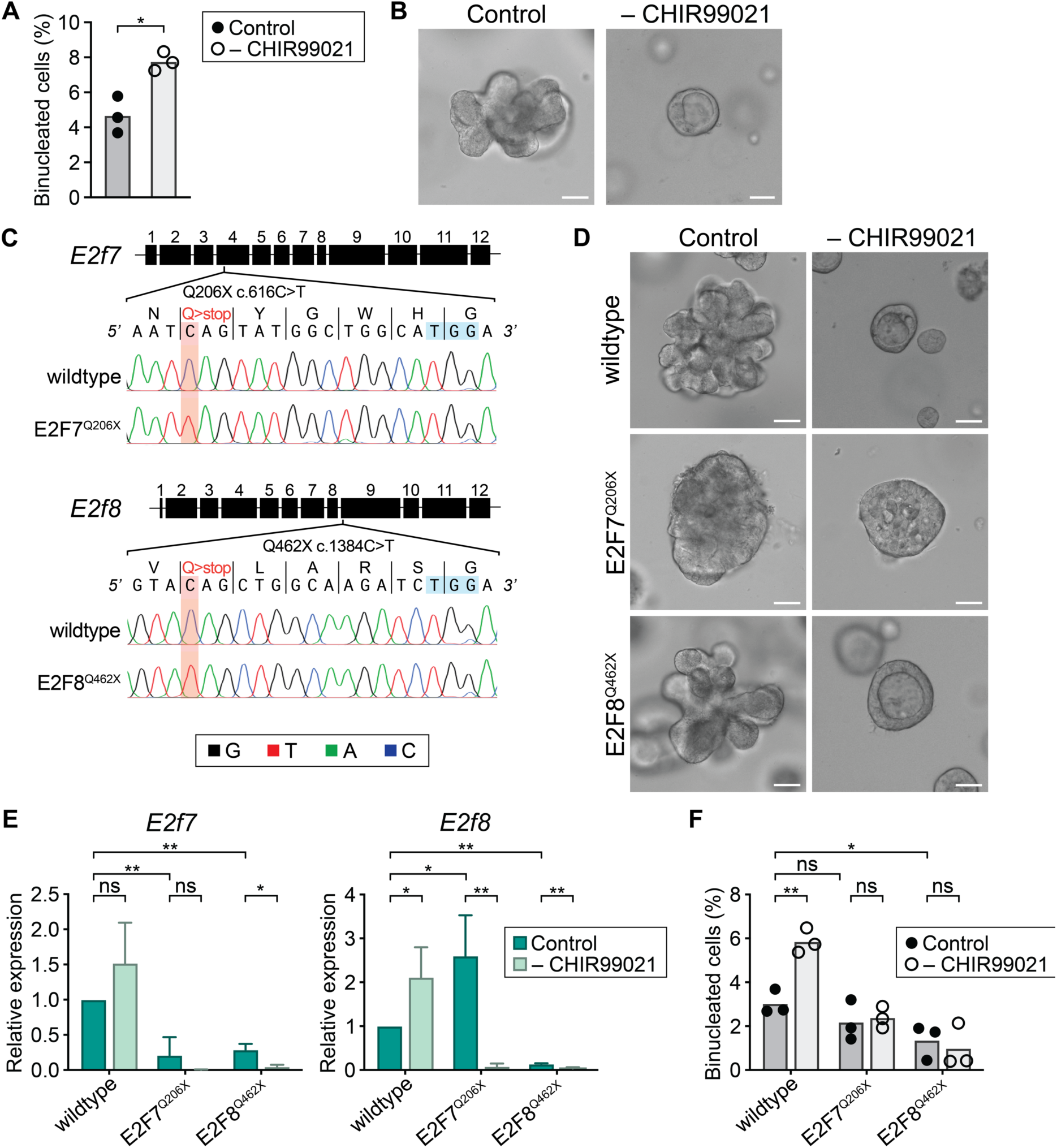
WNT signaling inhibits binucleation of human hepatocytes in an E2F7/8-dependent manner. **A**) Quantification of binucleated cells in Hep-Org 1 line with GFP-NLS and stained with CellMask Orange in control culture medium and after 3 days of CHIR99021 removal (n = 3 experiments, 200-300 cells analyzed per experiment, \**p*<0.05, Student’s *t*-test). **B**) Representative brightfield images of Hep-Org 1 in culture medium and upon CHIR99021 removal. Scale bars represent 50 µm**. C**) Base-editing strategy for introduction of premature stop codon in *E2f7* and *E2f8* open reading frames with sanger sequencing chromatographs of wildtype (top) or E2F7^Q206X^/E2F8^Q462X^ (bottom) lines confirming homozygous base changes (highlighted in red). PAM sequences are highlighted in blue. **D**) Representative brightfield images of wildtype, E2F7^Q206X^, and E2F8^Q462X^ Hep-Org 1 lines in culture medium and upon CHIR99021 removal. Scale bars represent 50 µm. **E**) Relative expression of *E2f7* or *E2f8* in wildtype and mutant lines as measured by RT-qPCR (n = 3 experiments, ns = not significant, \**p*<0.05, \*\**p*<0.01, Student’s *t*-test). Error bars represent standard deviation. **F**) Quantification of binucleated cells in wildtype and mutant lines expressing GFP-NLS and stained with CellMask Orange (n = 3 experiments, 200-300 cells analyzed per experiment, ns = not significant, \**p*<0.05, \*\**p*<0.01, Student’s *t*-test).

To determine if WNT-mediated repression of binucleation in Hep-Orgs is dependent on E2F7 and E2F8, as has been reported in mouse hepatocytes^21–23^, we used the CRISPR-based cytosine base editor (CBE) system^60,61^ to introduce premature stop codons in the endogenous *E2F7* and *E2F8* loci (**Fig. 4C**). The mutant lines showed differences in proliferation and morphology compared to wild-type Hep-Orgs: the E2F7^Q206X^ line proliferated slower and exhibited a dense morphology, while the E2F8^Q462X^ line grew at a comparable rate with a similar bunch-of-grapes morphology as the wildtype line (**Fig. 4D**). We confirmed by quantitative reverse transcription PCR (RT-qPCR) that the introduction of a premature stop codon resulted in nonsense-mediated decay (NMD) of the respective genes (**Fig. 4E**), and also found that the expression of *E2F8* is increased in the E2F7^Q206X^ line, while expression of *E2F7* is decreased in the E2F8^Q462X^ line. In normal culture conditions, the percentage of binucleated cells was lower in the E2F8^Q462X^ line than in wildtype, but was comparable between E2F7^Q206X^ and wildtype Hep-Org lines (**Fig. 4F**). Together, these suggest that knockdown of *E2F7* or *E2F8* individually is not sufficient to completely prevent binucleation, but that E2F7 and E2F8 function redundantly to promote binucleation.

Strikingly, we did not see an increase in the percentage of binucleated cells upon removal of CHIR99021 in the E2F7^Q206X^ and E2F8^Q462X^ mutant Hep-Org lines, suggesting that E2F7 and E2F8 are required to induce binucleation upon reduction of WNT signaling. (**Fig. 4F**). Interestingly, we found that although CHIR99021 removal in wildtype organoids leads to increased *E2F8* expression, the removal of CHIR99021 in E2F7^Q206X^ mutant organoids drastically reduces *E2F8* expression (**Fig. 4E**). Furthermore, E2F7^Q206X^ mutant organoids, in contrast to wild type and E2F8^Q462X^ mutant organoids, do not become round and cystic in the absence of CHIR99021(**Fig. 4D**). Taken together, our results suggest that E2F7 and E2F8 control hepatocyte differentiation and binucleation, and that similar to mouse hepatocytes, WNT signaling inhibits binucleation of human hepatocytes in an E2F7-and E2F8-dependent manner.

## Discussion

Although non-canonical cell cycles are crucial for the function of many organs, little is known on how they are initiated and executed in human cells, largely due to unavailability of suitable systems. In the present study, we used non-transformed, healthy human hepatocyte organoids to study how cells undergo non-canonical cell cycles and become binucleated. Our findings indicate that binucleated cells in Hep-Orgs arise by detachment of the cell membrane to the midbody in late cytokinesis. This late cytokinesis exit contrasts to endomitosis in rodent livers, where cells lack a central spindle and do not undergo cleavage furrow ingression^20–22^. Despite the differences in the mechanism by which cells inhibit cell division, we uncover a conserved function of WNT signaling along with E2F7 and E2F8 in the regulation of endomitosis, suggesting that similar mechanisms control the choice between canonical versus non-canonical cell cycles in rodents and human hepatocytes.

We find that on average, 3-15% of cells in the Hep-Org lines used in this study are binucleated. The differences in percentages of binucleated cells between the two Hep-Org lines likely reflects normal biological variation, as they are consistent with reported percentages of binucleated cells in fetal and neonatal human livers, ranging between 1.5-8%^30,31^. Notably, despite that the Hep-Org 2 line shows higher percentages of binucleated cells, this is not reflected in the percentage of cells undergoing endomitosis M phase, which is very similar between Hep-Org 1 and Hep-Org 2 lines. This may indicate that the higher percentage of binucleated cells in Hep-Org 2 is due to longer retention of binucleated cells instead of a higher rate of endomitosis. In both rodents and humans, the number of binucleated hepatocytes increases with age, with rodent livers containing on average around 80% binucleated cells^26^ and adult human livers consisting of up to 20% binucleated cells^30,31,62^. In mouse livers, the percentage of hepatocytes undergoing endomitosis increases drastically upon weaning, however it is unknown what triggers endomitosis cycles in human livers^21–23^. It is possible that changes in hepatic metabolism influence the choice between canonical and non-canonical M phases in human hepatocytes, but this remains to be determined.

From our live-imaging analyses of Hep-Orgs, we find that human hepatocytes undergoing endomitosis inhibit cell division during a late step in cytokinesis. Endomitotic hepatocytes display normal cytokinetic furrow ingression, but the ingressed membrane detaches from the midbody during late cytokinesis. We find that RacGAP1, Anillin, and CIT-K, which are involved in anchoring the midbody to the cell cortex, are present but lose their association to the cell membrane during endomitotic furrow regression. In addition, we find that CIT-K shows a distinct localization along the midbody arms in regressed structures. Interestingly, there is evidence that inhibition of CIT-K may also play a role in non-canonical cell cycles in the mouse brain^63^. Here, inhibition of CIT-K by Src kinase promotes the formation of polyploid neurons in the developing mouse neocortex^63^. It will be interesting to determine whether Src signaling is also involved in human hepatocyte endomitosis, as there is evidence that Src kinase activity promotes binucleation of mouse hepatocytes^64^.

Within the same cell type, endomitosis does not always occur in the same manner across different species. For instance, the majority of polyploid mouse cardiomyocytes are binucleated, having aborted cytokinesis at a late stage of M phase^65^, whereas the majority of polyploid human cardiomyocytes are mononucleated, and thus likely exit M phase before anaphase^66–70^. Our findings demonstrate that there are also differences in endomitosis between human and rodent hepatocytes^20–22^, raising the question whether cells evolved independent mechanisms to become polyploid, or whether there are evolutionarily conserved regulators of non-canonical cell cycles. In this study, we investigated the function of WNT signaling, which has previously been described to control non-canonical cell cycles in mice. Similar to murine hepatocytes, we find that WNT signaling inhibits human hepatocyte binucleation. Furthermore, we find that the atypical E2F family transcription factors E2F7 and E2F8 act downstream of WNT signaling to promote binucleation, demonstrating a conserved function of endomitosis regulators.

Interestingly, E2F7 and E2F8 have not only been implicated in endomitosis but also in endoreplication cycles in trophoblast giant cells^71–73^. As initiation of endoreplication requires inhibition of mitotic entry, it is possible that E2F7 and E2F8 act as general repressors for genes involved in M phase, including genes responsible for cytokinesis. Indeed, knockout of E2F7 and E2F8 in mouse livers leads to an increase in expression M-phase genes, and E2F8 has also been shown to bind to promoters of M-phase genes in human cells^23^. This raises the question of how the activity of E2F7 and E2F8 is specifically required to induce binucleation and does not lead to endoreplicative cycles in hepatocytes. One possibility is that in endomitotic cells, activation of E2F7 and E2F8 leads to transcriptional repression of all M-phase genes, but the available proteins remain stable at sufficient levels to allow mitotic entry but are not sufficient to complete cytokinesis. To test this hypothesis, one would require a detailed comparison of expression profiles between cells undergoing canonical and endomitosis M phases. However, several technical aspects currently render this challenging: firstly, we currently lack the knowledge to be able to predict whether a cell will undergo a canonical or endomitosis M phase, preventing isolation of this rare cell population; and secondly, M-phase regulators are expressed at low and fluctuating levels during the cell cycle, making it difficult to compare gene expression profiles between cells. Further dissection of the pathways that drive endomitosis may allow generation of mutant hepatocytes with higher levels of endomitosis, which will facilitate a detailed comparison between cells undergoing canonical and endomitosis M phase.

Taken together, our study provides insights into the mechanism by which human cells become binucleated through non-canonical cell cycles during tissue formation. We find that endomitosis in human hepatocytes occurs by loss of membrane anchorage to the midbody during M phase, and that binucleation is controlled by WNT signaling and E2F7/8 activities. As liver polyploidization is important to balance liver gene expression and proliferation^7,27,74^, understanding the modulation of non-canonical cell cycles may provide therapeutic opportunities in the treatment and prevention of diseases such as liver cancer.

## Supplementary Figure

**Figure S1.**
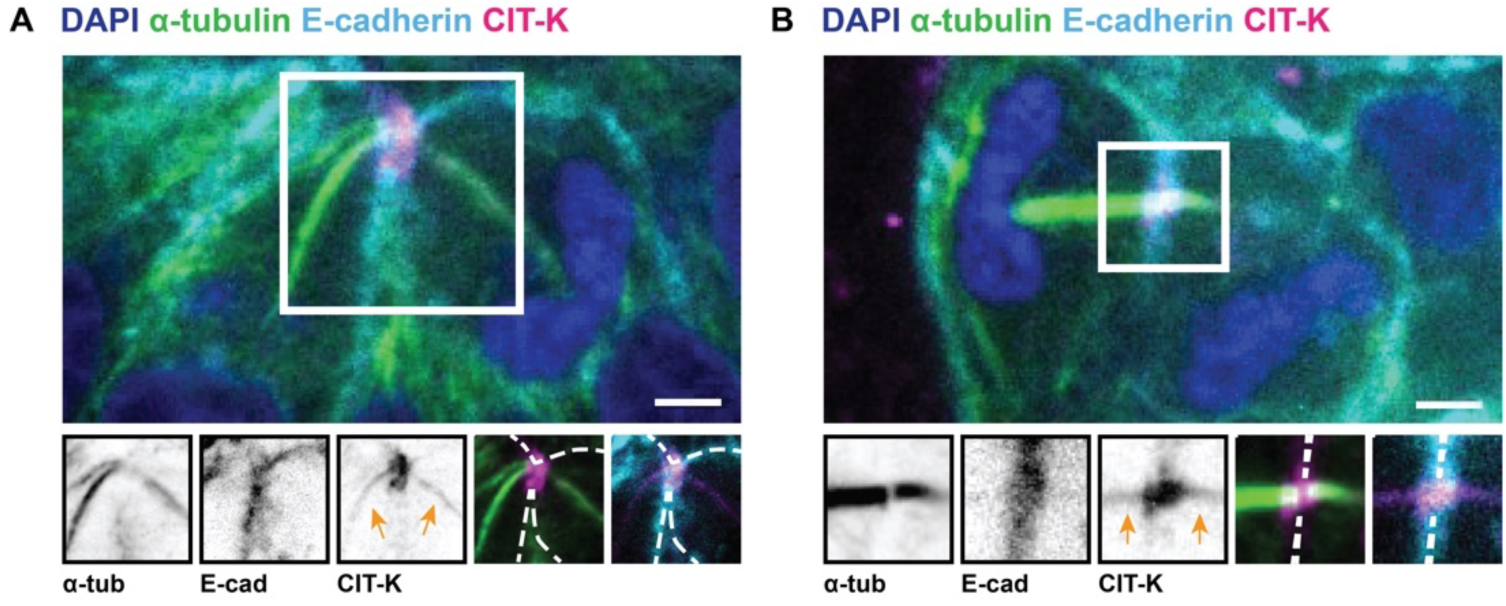
Representative stills of IF experiments showing atypical CIT-K co-localization with microtubules in early (A) and late telophase, ingressed (B). Scale bars represent 3 µm. Close-ups show single or double channel images of marked region of interest (white box). Orange arrows point to CIT-K signal in midbody arms. Dashed lines represent membrane outline. All images show one plane in the middle of the midbody.

## Materials and methods

### Organoid culture

Hep-Orgs used in this study were previously generated from human fetal livers that were obtained from Leiden University Medical Centre (MC) under approval of the Dutch Ethical Medical Council (Leiden University MC)^44,45^. All organoids were grown in Cultrex reduced growth factor basement membrane extract (BME), type 2 (R&D Systems, #3533-001) and cultured and passaged as previously described for Chol-Orgs^75^ and Hep-Orgs^44^. Culture medium components are listed in **Table 2**.

**Table 2.** Organoid culturing medium.

| Line | Component | Catalog number | Supplier | Concentration |
| --- | --- | --- | --- | --- |
| Chol-Org | Advanced DMEM/F12 | 12634 | Gibco |  |
|  | HEPES | 15630 | Gibco | 10 mM |
|  | GlutaMAX | 35050 | Gibco | 1x |
|  | Penicillin/Streptomycin | 15140 | Gibco | 100 U/mL |
|  | RSPO1 conditioned medium | - | (Made in house) | 10% |
|  | B27 minus vitamin A | 12587-010 | Thermo Scientific | 1x |
|  | Nicotinamide | 72340 | Sigma Aldrich | 10 mM |
|  | N-acetylcysteine | A7250 | Sigma Aldrich | 1.25 mM |
|  | FGF10 | 100-26 | Peprotech | 100 ng/mL |
| | A83-01 | 2939 | Tocris | 5 μM |
| | FSK | 1099 | Tocris | 10 μM |
|  | HGF | 100-39 | Peprotech | 25 ng/mL |
| | EGF | AF-100-15 | Peprotech | 50 ng/μL |
|  | Gastrin I | 3006 | Tocris | 10 nM |
| Hep-Org | Advanced DMEM/F12 | 12634 | Gibco |  |
|  | HEPES | 15630 | Gibco | 10 mM |
|  | GlutaMAX | 35050 | Gibco | 1x |
|  | Penicillin/Streptomycin | 15140 | Gibco | 100 U/mL |
|  | RSPO1 conditioned medium | - | (Made in house) | 15% |
|  | B27 minus vitamin A | 12587-010 | Thermo Scientific | 1x |
|  | Nicotinamide | 72340 | Sigma Aldrich | 10 mM |
|  | N-acetylcysteine | A7250 | Sigma Aldrich | 1.25 mM |
| | Y-27632 | Y0503 | Sigma Aldrich | 10 μM |
| | CHIR99021 | 4423 | Tocris | 3 μM |
|  | HGF | 100-39 | Peprotech | 50 ng/mL |
|  | FGF7 | 100-19 | Peprotech | 100 ng/mL |
|  | FGF10 | 100-26 | Peprotech | 100 ng/mL |
| | A83-01 | 2939 | Tocris | 2 μM |
|  | EGF | AF-100-15 | Peprotech | 50 ng/mL |
| | TGF-α | 100-16A | Peprotech | 20 ng/mL |
|  | Gastrin I | 3006 | Tocris | 10 nM |

### Generation of organoid lines

For lines expressing GFP-NLS, organoids were transduced with lentivirus containing pHR-sfGFP-NLS using the sandwich method adapted from 76. In short, organoids were dissociated using TrypLE (Gibco, #12605) to small clumps and plated on a layer of BME type 2 (R&D Systems, cat. no. 3533-001), incubated overnight in culture medium supplemented with 10 μg/mL polybrene (Santa Cruz Biotechnology, #sc-134220) containing concentrated lentiviral supernatant (Centriprep centrifugal unit - 10 kDa cutoff, Sigma Aldrich, #4305). The next day, the culture medium was replaced and a second layer of BME was added to transduced organoids to allow recovery, after which organoids were cultured normally. The generation of lines expressing endogenous Tubulin-mNeon and/or endogenous E-cadherin-tdTomato was performed using CRISPaint as previously described^45,77^. Premature stop codons in E2F7 and E2F8 were introduced in Hep-Org 1 line using CRISPR base editing as previously described^78^. All guide RNA sequences are listed in **Table 3**. Clonal organoid lines were obtained by selecting organoids grown from single cells following digestion with TrypLE. For genotyping, organoids were lysed at 65°C for 15 minutes followed by enzyme inactivation at 95°C for 5 minutes in lysis buffer containing 50 mM KCl (Avantor, #0509), 2.5 mM MgCl_2_ (Avantor, #0162), 10 mM Tris-HCl pH 8.3 (Invitrogen, #15504-020), 0.45% IGEPAL CA-630 (Sigma-Aldrich, #I8896), 0.45% Tween-20 (Sigma-Aldrich, #P1379), and 1 mg/mL Proteinase K (Sigma-Aldrich, #P2308), followed by PCR amplification using Q5 High-Fidelity DNA polymerase (New England Biolabs, #M0491) and Sanger sequencing (Macrogen Europe). Genotyping primers are listed in **Table 3**.

**Table 3.**
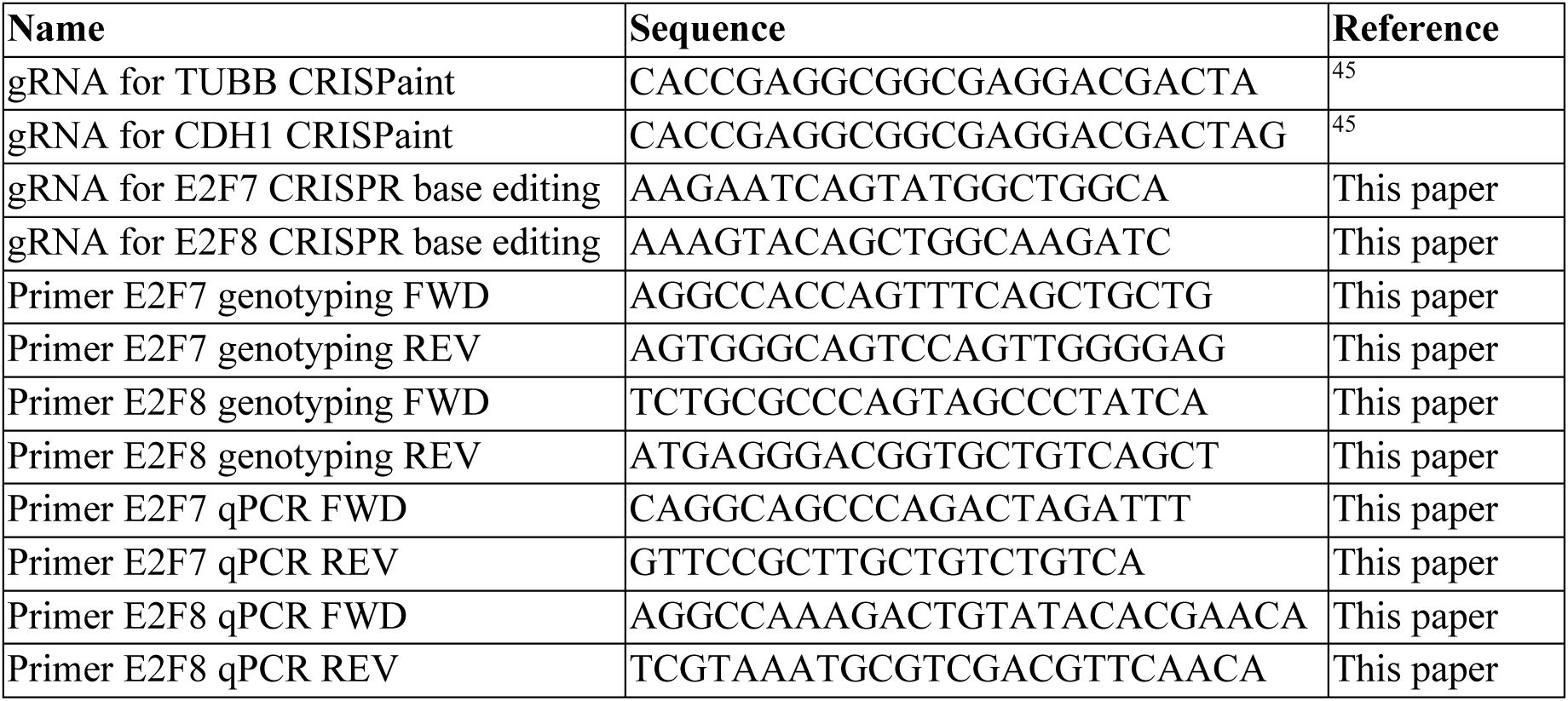
Oligo sequences.

### RNA isolation and RT-qPCR

RNA was extracted from organoids using TRIzol extraction (Invitrogen, #15596018). cDNA synthesis was performed using iScript cDNA synthesis kit (Bio-Rad, #1708891). qPCR was performed using iQ SYBR Green Supermix (Bio-Rad, #1708882) in a CFX96 Real-Time PCR Detection System (Bio-Rad). qPCR primers for E2F7 and E2F8 were designed using NCBI Primer-BLAST to target regions upstream of target base for CRISPR base editing and are listed in **Table 3**.

### Live imaging, immunofluorescence, and microscopy

For live-cell imaging, organoids were plated onto ibidi 8-well chambered coverslips with ibiTreat (ibidi, #80826) one day prior to imaging. Imaging was performed using a Nikon TiE microscope equipped with Borealis illumination unit (Andor), CSU-W1 spinning disk scanner unit (Yokogawa), and iXon-888 Ultra EMCCD camera (Andor) using a UPLSAPO-S 30x silicone objective (Olympus) with 1.5x optical zoom in an environmentally-controlled chamber held at 37°C with 5% CO_2_. Organoids were imaged for 24-72 hours with 5-minute intervals, with a total of 12 z-slices spaced 4 µm apart per position.

To quantify the number of binucleated cells, Hep-Org expressing GFP-NLS were grown on ibidi 8-well chambered coverslips (ibidi, #80826) and incubated with CellMask Orange (1:2,000, Invitrogen, #C10045) for 20 minutes, after which the staining solution was removed, and fresh culture medium was added. Organoids were imaged immediately using the same spinning disk microscope system as described above.

For immunofluorescent staining, organoids were plated onto ibidi 8-well chambered coverslips 3 days prior to fixation. Organoids were fixed in the BME droplets in 4% paraformaldehyde in PBS (Electron Microscopy Sciences, #RT15710) for 10 minutes, followed by permeabilization and blocking in blocking buffer containing 1% bovine serum albumin (Sigma-Aldrich, #A3294), 10% DMSO (Sigma-Aldrich, #D8418), and 2% Triton-X100 (Sigma-Aldrich, #T8787) in PBS. Organoids were incubated overnight at 4°C with rat anti-α-tubulin (1:1,000, Novus Biologicals, #NB600-506), goat anti-RacGAP1 (1:1,000, Abcam, #ab2270), mouse anti-anillin (1:500, Sigma Aldrich, #MABT96), or mouse anti-CIT-K (1:500, BD Biosciences, #611376) in blocking buffer. Organoids were washed in PBS followed by overnight incubation with species-matched secondary antibodies conjugated to Alexa fluor dyes (Invitrogen), after which samples were washed thoroughly with PBS and co-stained with DAPI (1:1,000, Sigma-Aldrich, #32670). Organoids were rinsed in PBS and imaged in PBS using a Plan Apo λ 60x oil objective (Nikon, #MRD01605) in Nikon Ti2 microscope equipped with L6Cc illumination unit (Oxxius), CSU-X1 spinning disk scanner unit (Yokogawa), with a C11440-22C camera (Hamamatsu).

For overview images of organoid mutant lines, brightfield images were taken on EVOS FL microscope (Invitrogen, #AMF4300).

#### Data analysis

All images were processed and analyzed using FIJI79. In our analyses, only organoids containing upwards of 10 cells were included. For quantification of percentages of binucleated cells, only healthy-looking cells with round nuclei were included in the analyses. For analysis of M phase types, the Mitotic Scoring Tool macro80 was used to aid M phase annotation.

##### Acknowledgements

We thank H. Hu, B. Artegiani, D. Hendriks, M. Geurts, and S. van den Brink (Hubrecht Institute, Utrecht, the Netherlands) for organoids, reagents, advice, and technical assistance, A. de Graaff and P. Toonen from the Hubrecht Imaging Centre for microscopy support, D. Bodor for the FIJI macro for M phase quantification, and G. Kops and M. Gloerich for advice throughout the project and comments on the manuscript. This work was supported by funding from the Cancer Genomics Center (CGC.nl).

